# Proximate causes and consequences of intergenerational influences of salient sensory experience

**DOI:** 10.1101/509596

**Authors:** Hadj S Aoued, Soma Sannigrahi, Sarah C Hunter, Nandini Doshi, Anthony Chan, Hasse Walum, Brian G Dias

**Affiliations:** Division of Behavioral Neuroscience and Psychiatric Disorders – Yerkes National Primate Research Center – Atlanta GA; Division of Neuropharmacology and Neurologic Diseases – Yerkes National Primate Research Center – Atlanta GA; Department of Human Genetics – Emory University School of Medicine – Atlanta GA; Division of Microbiology and Immunology – Yerkes National Primate Research Center – Atlanta GA; Silvio O. Conte Center for Oxytocin and Social Cognition, Center for Translational Social Neuroscience, Yerkes National Primate Research Center, Emory University, Atlanta, GA, USA; Department of Psychiatry and Behavioral Sciences – Emory University School of Medicine – Atlanta GA

## Abstract

Salient sensory environments experienced by a parental generation can exert intergenerational influences on offspring, including offspring not conceived at the time of the parental experience. While these data provide an exciting new perspective on biological inheritance, questions remain about causes and consequences of intergenerational influences of salient sensory experience. We have previously shown that exposing male mice to a salient olfactory experience like olfactory fear conditioning results in offspring demonstrating a sensitivity to the odor used to condition the paternal generation and possessing an enhanced neuroanatomical representation for that odor. In this study, we first injected RNA extracted from sperm of male mice that underwent olfactory fear conditioning into naïve single cell zygotes and found that both male and female adults that develop from these embryos have increased sensitivity and enhanced neuroanatomical representation for the odor (Odor A) with which the paternal male had been conditioned. Next, we found that female, but not male offspring sired by males conditioned with Odor A show enhanced freezing when presented with Odor A after being exposed to a sub-threshold olfactory fear conditioning protocol that consisted of only a single Odor A + shock pairing. Our data provide evidence that RNA found in the paternal germline after exposure to salient sensory experiences can contribute to intergenerational influences of such experiences, and that such intergenerational influences confer an element of adaptation to the filial generation. In so doing, our work suggests that some causes (sperm RNA) and consequences (behavioral flexibility) of intergenerational influences of parental experiences are conserved across experiences as diverse as stressors, dietary manipulations, and sensory experiences.

## INTRODUCTION

Sensory cues abound in the umwelt of an organism. Appropriate responses to physical and chemical stimuli in the environment are crucial for an organism to function optimally and survive. While such responses are influenced by experiences that come to be associated with these stimuli, accumulating evidence suggests that the legacy of parental experiences with stimuli can significantly shape responses in future generations. Organisms mount defense responses to physical and chemical cues from predators, but the development of such responses can arise *de novo* during development as a consequence of parental exposure to these same predators [1, 2]. Environmental insults like droughts, endocrine disruptors and dietary manipulations influence responses of offspring in both plants and animals even with the offspring not directly experiencing the insults [3–7]. Also, offspring of a variety of species ranging from humans to mice exhibit a preference for the odors of foods consumed by mothers during pregnancy or lactation [8–10]. These data are representative of a burgeoning field of research that suggests the profound influence that salient sensory environments exert on future generations. However, despite an appreciation for this phenomenon of intergenerational influences of salient sensory events, very little is known about how information about salient sensory cues is inherited via the parental gametes and the consequences of this inheritance on the physiology and behavior of offspring.

The olfactory system is an ideal system to successfully address how experience with a salient environmental cue can exert influences across generations [11–19]. Training mice to associate a specific odor (e.g. Odor A) with a mild foot-shock results in these mice developing fearful behavior toward Odor A and possessing increased neuroanatomical expression of odorant receptors that detect Odor A [20, 21]. From an intergenerational perspective, we have demonstrated that after mice have been exposed to Odor A + shock conditioning their naïve offspring possess a sensitivity to Odor A and increased neuroanatomical expression of odorant receptors that detect Odor A [22, 23]. Ours are not the only studies to have reported intergenerational imprints of parental olfactory experience. Olfactory conditioning of fruit-flies influences the behavior of offspring [24]. Appetitive and adverse maternal experience with odors in rodents is transmitted to neonatal pups thereby impacting the neuroanatomy, physiology and behavior of the pups when they detect that odor [10, 12]. Odor-based learning has been shown to have intergenerational effects in offspring [25]. Building on our studies using olfactory fear conditioning in a parental generation of mice and examining influences in offspring, we ask how intergenerational influences of salient olfactory experiences can be inherited across generations and what consequence such influences have on the behavior of the offspring as they navigate their own umwelts.

With RNA found in sperm of male rodents shown to be mediators of intergenerational influences of stress and dietary manipulations [26–30], we examined whether RNA found in the sperm of male mice exposed to olfactory fear conditioning played any influence in intergenerational influences of salient olfactory experience. To do so, we injected RNA obtained from sperm of male mice exposed to olfactory fear conditioning with Odor A into naïve single cell zygotes and measured olfactory sensitivity and olfactory neuroanatomy related to Odor A in adulthood. While intergenerational influences of salient parental environments have traditionally been thought to constrain biology in offspring, there is a growing body of evidence to suggest that such influences can afford behavioral flexibility to offspring and be adaptive. To examine this possibility in the context of salient sensory experiences, we exposed male mice to olfactory fear conditioning with Odor A and then examined learning and memory in offspring after conditioning them to Odor A.

## MATERIALS & METHODS

### Animals

Experiments were conducted with 2-3 month old sexually and odor-inexperienced animals. C57BL/6J animals, M71-LacZ animals maintained in mixed 129/Sv x C57Bl/6J background, and MOR23-GFP animals maintained in mixed 129/Sv x C57Bl/6J background were bred in the Yerkes Neuroscience animal facility. Animals were housed on a 12 h light/dark cycle in standard groups cages (5/cage) with ad libitum access to food and water, with all experiments conducted during the light half of the cycle. All procedures were approved by the Institutional Animal Care and Use Committee of Emory University, and followed guidelines set by the National Institute of Health.

### Experiments related to determining causes of intergenerational influences of salient olfactory experience (Fig. 1)

#### Olfactory treatment of parental F0 generation

**F0-Trained** – Males were trained to associate Acetophenone or Lyral presentation with mild foot-shocks. For this purpose, the Startle-Response system (SR-LAB, San Diego Instruments) was modified to deliver discrete odor stimuli as previously described. The animals were trained on 3 consecutive days with each training day consisting of 5 trials of odor presentation for 10 secs co-terminating with a 0.25 sec 0.4mA foot-shock with an average inter-trial interval of 120 secs. Both Acetophenone and Lyral (from Sigma, and IFF, respectively) were used at a 10% concentration diluted with Propylene Glycol. These odors were chosen based on prior work that demonstrated that the M71 odorant receptor is activated by Acetophenone, and that the MOR23 odorant receptor is activated by Lyral [14, 19, 31]. **F0-Exposed** – Males were treated like the F0-Trained group with the main important exception that odor presentations were not accompanied by any foot-shocks.

**Fig. 1:**
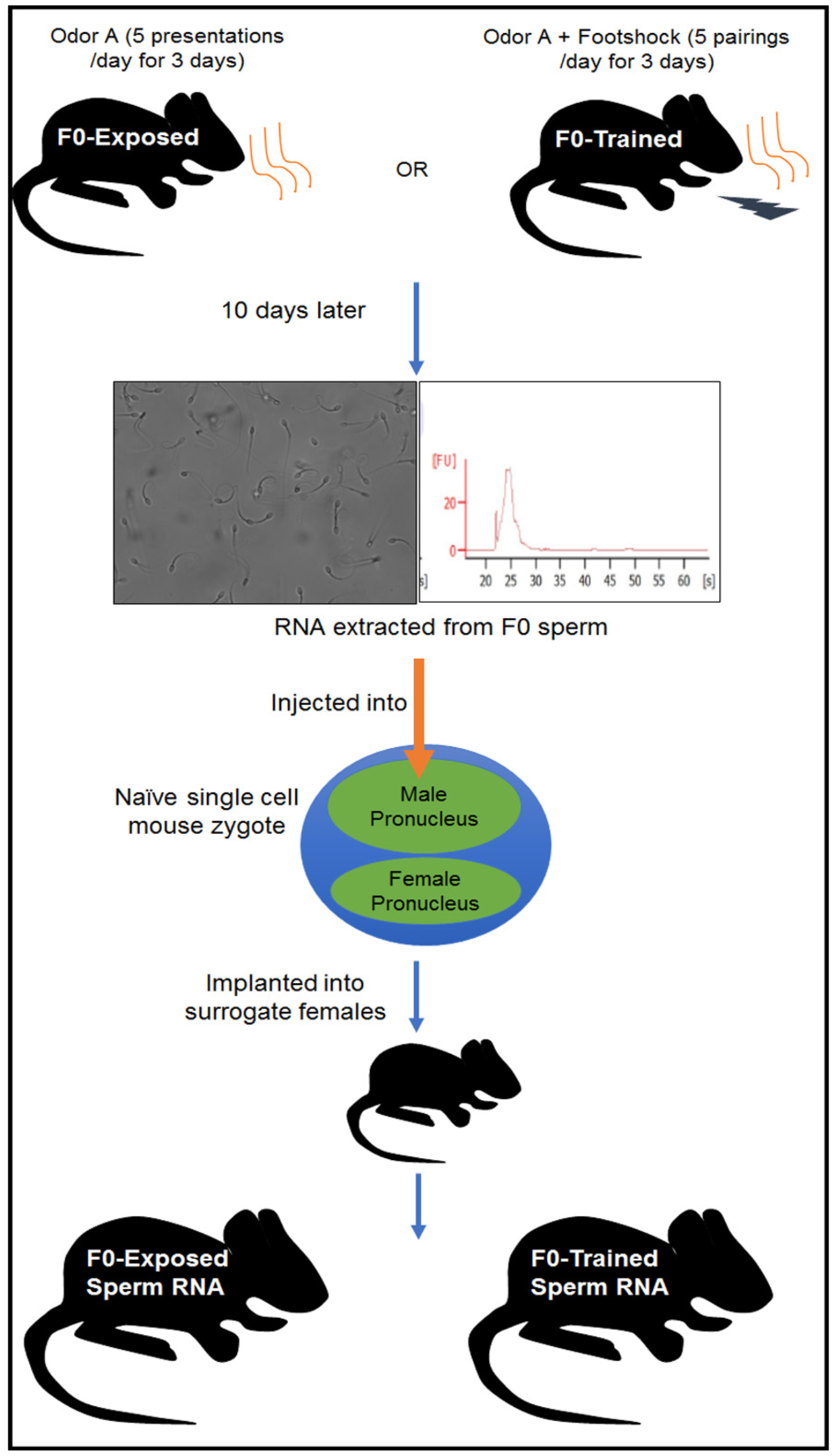
Experimental design to determine the contribution of sperm RNA to behavioral and neuroanatomical influences of salient paternal olfactory experiences across generations. Male mice were either exposed to an odor (e.g. Odor A) (F0-Exposed group) or exposed to Odor A while mild foot-shocks were paired with Odor A presentations (F0-Trained group). Ten days later, pure sperm were collected post-mortem and RNA extracted from these sperm. These RNA from sperm of F0-Exposed or F0-Trained males were then injected into the male pronucleus of naïve single cell mouse zygotes. The zygotes were implanted into surrogate female mice and experiments were performed on them at 2-months of age.

#### Sperm collection from F0 males and RNA extraction from sperm

Sperm was collected from F0-Exposed or F0-Trained mice that were treated with either Acetophenone or Lyral. Mice were anesthetized with isoflurane then euthanized by cervical dislocation. The abdominal cavity was opened using surgical scissors and the cauda epididymis was removed then transferred into a well of a 24-well plate that contained 500 ul of 1x PBS at room temperature. The cauda epididymis was punctured to allow the sperm to be released, using a 23G needle and a pair of forceps. The 24 well plate was placed into a 37C incubator with 5% CO2 for one hour. A gradient procedure using a Pure Sperm kit (Nidacon cat# PS40-100, PS80-100, PSW-100) was conducted at room temperature in order to obtain a pure sperm cell pellet. Using a pipette with sterile tip, 2 ml of pure sperm 80 was added into a 15 mL conical centrifuge tube. Then, with a new sterile pipette tip, 2 ml of pure sperm 40 was carefully layered on the top of pure sperm 80 layer. The cauda epididymis was removed from the solution and the liquefied semen was carefully layered onto the pure sperm gradient. Tubes were centrifuged at 300g for 20 minutes to allow the separation of the sperm cells from the rest of the tissue. A new sterile pipette was used in circular movement from the surface to aspirate everything except the pellet and 4-6 mm of pure sperm 80. With a new sterile pipette, the sperm pellet was transferred to a new tube. The pellet was resuspended in 5 ml of pure sperm wash buffer. The tubes were centrifuged at 500g for 10 minutes then all the fluid was aspirated and discarded except the pure sperm pellet. Sperm from three mice of the same F0 treatment condition were pooled. Purity of sperm was confirmed under a microscope. The pooled sperm pellet was stored in −80°C until used for RNA extraction. RNA was extracted from sperm that was collected following the TRIzol procedure (Invitrogen, Life Technologies). The concentration was measured using Qbit and adjusted to 1ng/ul before injection into zygote.

#### Isolation of zygote and RNA microinjection

To stimulate follicular growth, 2-3 month old behaviorally naive female mice were superovulated by first receiving an intraperitoneal injection of pregnant mare’s serum gonadotropin (PMSG; 5-10 IU) and then human Chorionic Gonadotropin (hCG; 5-10 IU), 48 hours after PMSG injection. Females were placed with behaviorally naïve male breeders overnight and vaginal plug was examined the following morning to confirm successful intercourse. Single cell zygotes were collected post-mortem for microinjection of RNA contained in sperm of treated F0 males. The zygotes were cultured in potassium-supplemented simplex optimised medium (KSOM, EMD Millipore) at 37°C in 5% CO2 incubator and then injected using a microinjector setup (XenoWorks Micromanipulator and Digital Microinjector by Shutter Instruments). The RNA was injected into the male pronucleus. The swelling of the pronucleus was used as an indication of a successful microinjection. The zygotes were cultured overnight in KSOM (EMD Millipore) covered with mineral oil and incubated in an incubator maintained at 37°C and with 5% CO2. Embryo development was accessed one day later by visualizing whether the zygotes reached the 2-cell stage. Genotype of the zygotes was M71-LacZ/+;MOR23-GFP/+.

#### Vasectomizing male mice, generating pseudo-pregnant females for embryo transfer, and embryo transfer into surrogate females

Two-month old ICR behaviorally naive male mice were sterilized. Briefly, vas deferens that is associated with the testis was identified and approximately 1cm of the segment was removed using sterile surgical technique followed by cauterization. After two weeks of recovery, sterility of the males was assessed by mating with naïve females. Only sterile males were used to generate pseudo-pregnant females as surrogate for embryo transfer. Briefly, 2-5 months old ICR surrogate behaviorally naive females were randomly selected from a cohort of adult females and placed into the cage of the sterile male overnight on the same day when RNA was injected to the single-cell zygotes. Pseudopregnancy was assessed by vaginal plug on the next day. Two-cell embryos (10-20 in number) were unilaterally transferred into the oviduct of a pseudo-pregnant female.

#### Odor-Potentiated Startle of injected zygotes as in adulthood (F1 offspring)

We measured baseline behavioral sensitivity of adult mice derived from RNA-injected zygotes to odors using an Odor Potentiated Startle (OPS) behavioral assay that we and others have used previously [22, 23, 32] and that measures acoustic startle response to a noise burst. Animals were habituated to the startle chambers for 5-10 minutes on 3 separate days. On the day of testing, animals were first exposed to 15 Startle-alone (105 dB noise burst) trials (Leaders), before being presented with 10 Odor+Startle trials randomly intermingled with 10 Startle-alone trials. The Odor+Startle trials consisted of a 10 sec odor presentation co-terminating with a 50 msec 105 db noise burst. For each animal, an Odor-Potentiated Startle (OPS) score was computed by subtracting the startle response in the first Odor+Startle trial from the startle response in the last Startle-alone Leader. This OPS score was then divided by the last Startle alone leader and multiplied by 100 to yield the percent OPS score (% OPS) reported in the results. Just like prior interpretations of this assay, we view higher % OPS readings as an indication of an increased sensitivity to the odor tested.

#### Neuroanatomy on F1 olfactory bulb and MOE

##### β-galactosidase staining

Brains were rapidly dissected and placed into 4% paraformaldehyde for 10 minutes at room temperature, after which they were washed 3 times in 1X PBS for 5 minutes each time. M71-LacZ was stained for β-galactosidase, using 45 mg of X-gal (1 mg/ml) dissolved in 600 ul of DMSO and 45 ml of a solution of 5 mM potassium ferricyanide, 5 mM potassium ferrocyanide, and 2 mM MgCl in 1X PBS, and incubated at 37°C for 3 hours.

##### Measurement of glomerular area in the olfactory bulb

A microscope-mounted digital camera was used to capture high-resolution images of the β-galactosidase stained M71 glomeruli at 40X magnification. Images were converted to grayscale and equalized for background brightness. The distribution of pixel brightness was exported in ImageJ as gray levels from 0 = black to 255 = white. X-gal labeled glomerular area was quantified as pixels, less than a set threshold gray level of 150 (optimized for axon vs background). Each glomerulus was traced using the lasso tool in Photoshop and the area was recorded from the histogram tool.

##### Western Blotting

10uL of protease cocktail inhibitor mix from Sigma (Cat# P8340) was mixed with 1mL and 500 uL of this mixture was added to a tube containing a fresh frozen MOE. The MOE was homogenized in this mixture using a plastic pestle and then an electric homogenizer. The tissue was then placed on a shaker for 15 min at 4°C to ensure complete lysis. The lysate was centrifuged at 4°C for 10 minutes at 12,000 rpm. The supernatant was transferred to a new tube on ice. Amount of protein in the lysate was determined using the Pierce BCA Protein Assay Kit. Standard SDS-PAGE was performed using 15-40 ug of protein per sample. After gel electrophoresis and transfer of the protein to Nitrocellulose membrane, the blots were probed with primary antibody (details noted below), the antibody detected by a peroxidase-coupled secondary antibody and signal detected using ECL substrate (Super Signal West Dura Extended Duration Substrate) and a BioRad Chemidoc MP-Imaging system.

##### Detecting LacZ

Primary Antibody – mouse anti-LacZ (Cat # 40-1a) from Developmental studies Hybridoma Bank; 1:200 dilutions with 2.5% nonfat dry milk in 1X TBS-0.1% Tween20, overnight on a rocker at 4 °C. Secondary Antibody – Anti mouse HRP, Cat # 7076S (Cell Signaling); 1:2000 dilution with 2.5% nonfat dry milk in 1X TBS-0.1% Tween20, 1 hour at room temperature.

##### Detecting GFP

Primary Antibody – Rabbit anti-GFP (ab6556) from Abcam; 1:2500 dilutions with 2.5% nonfat dry milk in 1X TBS-0.1% Tween20, overnight on a rocker at 4 °C. Secondary Antibody – Anti rabbit HRP; Cat # 7074S (Cell Signaling) 1:2000 dilution with 2.5% nonfat dry milk in 1X TBS-0.1% tween 20, 1 hour at room temperature.

##### Detecting β-actin as a loading control

Primary antibody – anti-Beta Actin (8H10D10) from Cell Signaling technology, dilution 1:5000 with 2.5% nonfat dry milk in 1X TBS-0.1% Tween20, 1 hour at room temperature. Secondary Antibody – Secondary Antibody: Anti mouse HRP, Cat # 7076S (Cell Signaling); 1:2000 dilution with 2.5% nonfat dry milk in 1X TBS-0.1% Tween20, 1 hour at room temperature. The amount of protein in every sample was quantified relative to β-actin by performing optical density measurements using ImageJ.

### Experiments related to determining consequences of intergenerational influences of salient olfactory experience (Fig. 2)

#### Olfactory treatment of parental F0 generation

C57BL/6J males were treated as noted above. Briefly, **F0-Trained** males were trained to associate Acetophenone or Lyral presentation with mild foot-shocks. The animals were trained on 3 consecutive days with each training day consisting of 5 trials of odor presentation for 10 secs co-terminating with a 0.25 sec 0.4mA foot-shock with an average inter-trial interval of 120 secs. **F0-Exposed** males were treated like the F0-Trained group with the main important exception that odor presentations were not accompanied by any foot-shocks. Ten days later, each mouse from these two groups was mated with naïve C57BL/6J female mice. Thirteen days later, animals were separated in order to prevent any interaction between the males and their offspring.

**Fig. 2:**
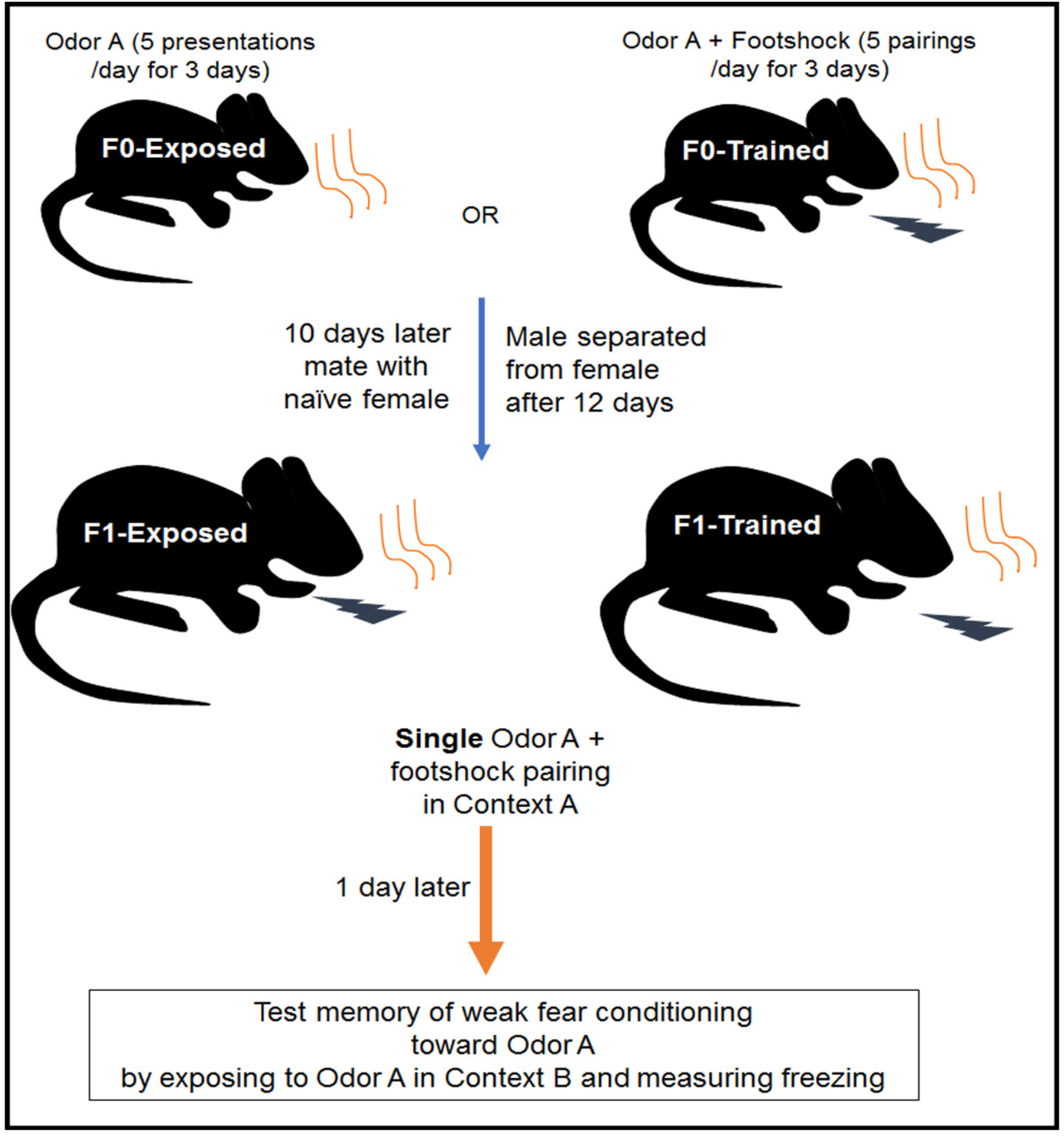
Experimental design to determine the consequence of paternal olfactory experience on olfactory learning in offspring. Male mice were either exposed to an odor (e.g. Odor A) (F0-Exposed group) or exposed to Odor A while mild foot-shocks were paired with Odor A presentations (F0-Trained group). Ten days later, these males were mated to naïve female mice, separated from the females after 12 days and experiments were performed on F1 offspring at 2-months of age. F1 animals were exposed to single trial olfactory conditioning in Context A (Odor A + foot-shock) and memory of this conditioning was measured one day later by presenting Odor A in Context B.

#### Testing for altered threshold for learning in the F1 generation

F1 generation adult mice were habituated for one day to the San Diego Instruments testing chambers. One day later, mice were trained to associate a single 10 sec Acetophenone or Lyral odor presentation with a co-terminating 0.25 sec 0.6mA foot-shock. Both Acetophenone and Lyral (from Sigma, and IFF, respectively) were used at a 10% concentration diluted with Propylene Glycol. The next day, animals were habituated to a completely different context, Mouse Habitest Chambers (Coulbourn Instruments). Finally, one day later in the Mouse Habitest Chambers animals were exposed to seven minutes of air followed by a 30-second presentation of Acetophenone or Lyral, and another seven minutes of air. As a measure of learning and consolidating memory about the single-trial olfactory fear conditioning, freezing behavior prior and in response to the odor presentation was recorded and analyzed using the FreezeFrame software. Our pilot experiments demonstrated that the single-trial olfactory fear conditioning was a weak conditioning protocol that resulted in baseline levels of freezing to the odor during the testing session allowing us to examine any augmentation of learning and memory as a consequence of a pre-existing sensitivity to the F0 conditioning odor.

#### Statistics

GraphPad Prism and the R package “nlme” was used to conduct statistical analyses. Data were analyzed using unpaired t-tests. For all experiments where potential litter effects (nonindependence of observations from the same litter) could arise, we ran mixed effect models including litter as a random effect [33] (random intercept) using the R package “nlme”, and statistics from these models are reported. All results are presented as Mean ± SEM, with *p<0.05, **p<0.01, ***p<0.001 and ****p<0.0001 as measures of significance and Cohen’s D is reported as an index of effect size. Effect sizes for experiments including animals from the same litter were calculated by taking the average per litter (to avoid issues due to correlated observations) and using these averages to estimate Cohen’s D. Although some statistical tests in which litter effects were accounted for did not reach significance, we observed consistently large effects sizes across experiments (and significance when standard unpaired t-tests were used to analyze the data). This could mean that statistical power, rather than the absence of underlying effects, explains why in some cases statistical significance was not observed.

## RESULTS

### Injection of RNA found in sperm of F0-Odor A-Trained males into naïve zygotes increases behavioral sensitivity to Odor A in adulthood

F0-Ace treatment: Injecting RNA extracted from sperm of F0-Ace-Trained-M71LacZ males into naïve single cell zygotes increased behavioral sensitivity of female and male animals to Acetophenone in adulthood (F0-Ace Trained Sperm RNA) compared to the behavioral sensitivity to Acetophenone measured in female and male animals that developed from zygotes injected with RNA extracted from sperm of F0-Ace-Exposed-M71LacZ males (F0-Ace Exposed Sperm RNA). (Female data: t=2.34, df = 10, p< 0.05, Cohen’s D = 1.4. Male data: t=3.33, df = 11, p< 0.01, Cohen’s D = 1.63) (Fig. 3A,B).

**Fig. 3:**
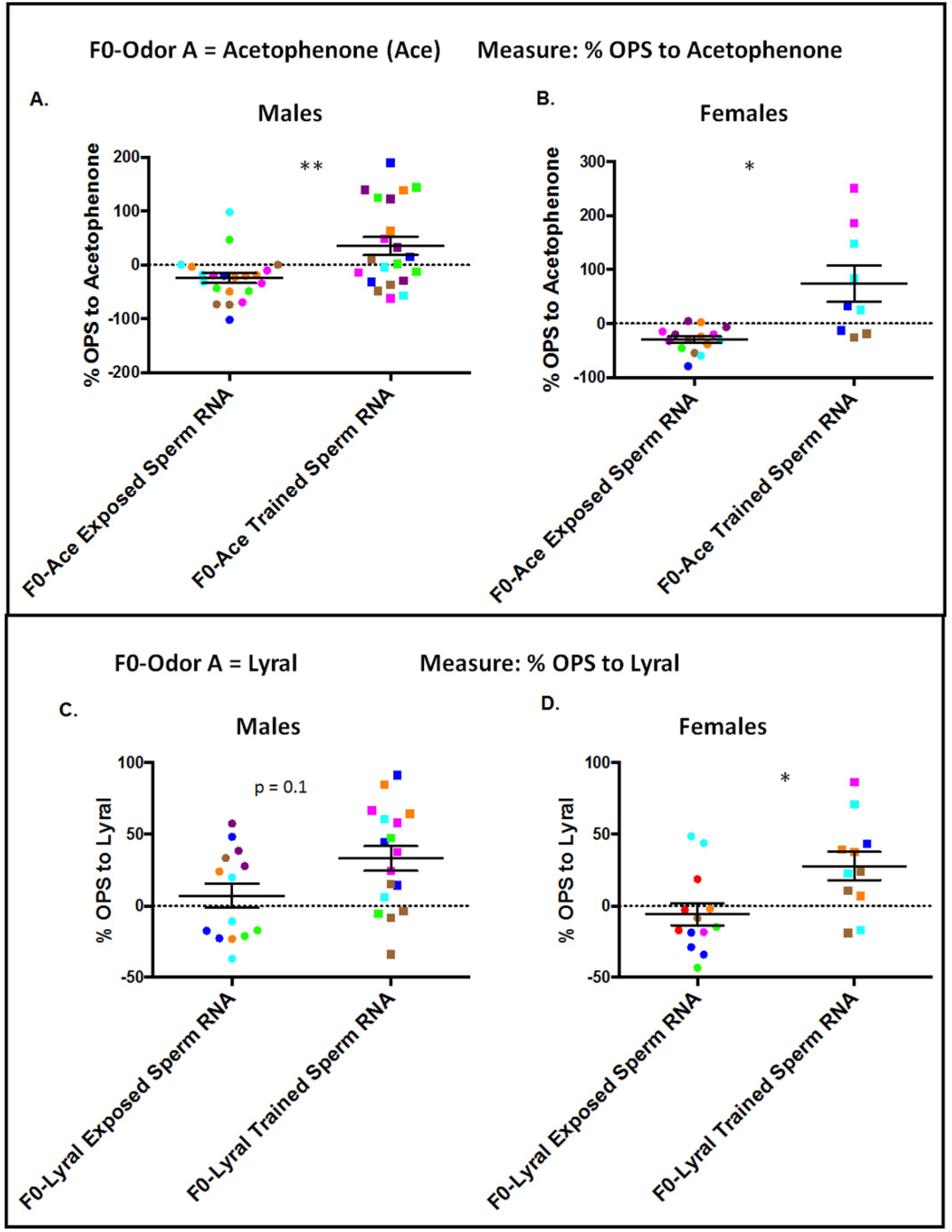
Sperm RNA contribute to behavioral imprints of salient olfactory experiences across generations. (A,B) Sensitivity to Acetophenone (Ace) after zygotes injected with RNA from sperm of F0 males that were treated with Ace. Male and female mice that developed from zygotes into which RNA from sperm of F0-Ace Trained males had been injected (F0-Ace Trained Sperm RNA, Males: n = 21, 7 litters, Females: n = 9, 5 litters) have a higher sensitivity to Acetophenone compared to mice that developed from zygotes into which RNA from sperm of F0-Ace Exposed males had been injected (F0-Ace Exposed Sperm RNA, Males: n = 21, 6 litters, Females: n = 15, 7 litters). **(C,D) Sensitivity to Lyral after zygotes injected with RNA from sperm of F0 males that were treated with Lyral.** Male and female mice that developed from zygotes into which RNA from sperm of F0-Lyral Trained males had been injected (F0-Lyral Trained Sperm RNA, Males: n = 17, 6 litters, Females: n = 11, 5 litters) have a higher sensitivity to Lyral compared to mice that developed from zygotes into which RNA from sperm of F0-Lyral Exposed males had been injected (F0-Lyral Exposed Sperm RNA, Males: n = 14, 6 litters, Females: n = 13, 7 litters). Data presented as Mean ± SEM. Colors denote animals from a litter. * p < 0.05, ** p < 0.01.

F0-Lyral treatment: Injecting RNA extracted from sperm of F0-Lyral-Trained-MOR23GFP males into naïve single cell zygotes increased behavioral sensitivity of female and male animals to Lyral in adulthood (F0-Lyral Trained Sperm RNA) compared to the behavioral sensitivity to Lyral measured in female and male animals that developed from zygotes injected with RNA extracted from sperm of F0-Lyral-Exposed-MOR23GFP males (F0-Lyral Exposed Sperm RNA). (Female data: t=2.41, df = 10, p< 0.05, Cohen’s D = 1.57. Male data: t=1.79, df = 10, p = 0.104, Cohen’s D = 1.07) (Fig. 3C,D).

### Injection of RNA found in sperm of F0-OdorA-Trained males into naïve zygotes enhances representation of Odor A-related neuroanatomy in the adult olfactory system

#### F0-Ace treatment

Adult female and male mice that developed from zygotes injected with RNA extracted from sperm of F0-Ace-Trained-M71LacZ males (F0-Ace Trained Sperm RNA) had larger M71-LacZ glomeruli in adulthood compared to adult mice that developed from zygotes injected with RNA extracted from sperm of F0-Ace-Exposed-M71LacZ males (F0-Ace Exposed Sperm RNA). (Female data: Dorsal glomeruli - t=1.62, df = 9, p=0.1387, Cohen’s D = 0.95. Medial glomeruli - t=4.04, df = 8, p< 0.01, Cohen’s D = 2.51. Male data: Dorsal glomeruli - t=1.88, df = 10, p=0.08, Cohen’s D = 1.01. Medial glomeruli - t=3.68, df = 7, p< 0.01, Cohen’s D = 1.81) (Fig. 4A-F). To ensure another measure of OR expression in the MOE of adult mice developing from zygotes that had been injected with RNA extracted from sperm of Acetophenone-treated F0 males, we performed western blotting on M71-LacZ MOE with a LacZ antibody that we have previously validated for its efficacy in detecting LacZ staining. This approach revealed that F0-Ace Trained Sperm RNA animals had higher LacZ expression in the MOE than F0-Ace Exposed Sperm RNA (t=3.144, df = 9, p< 0.05, Cohen’s D = 1.9) (Fig. 4G,H).

**Fig. 4:**
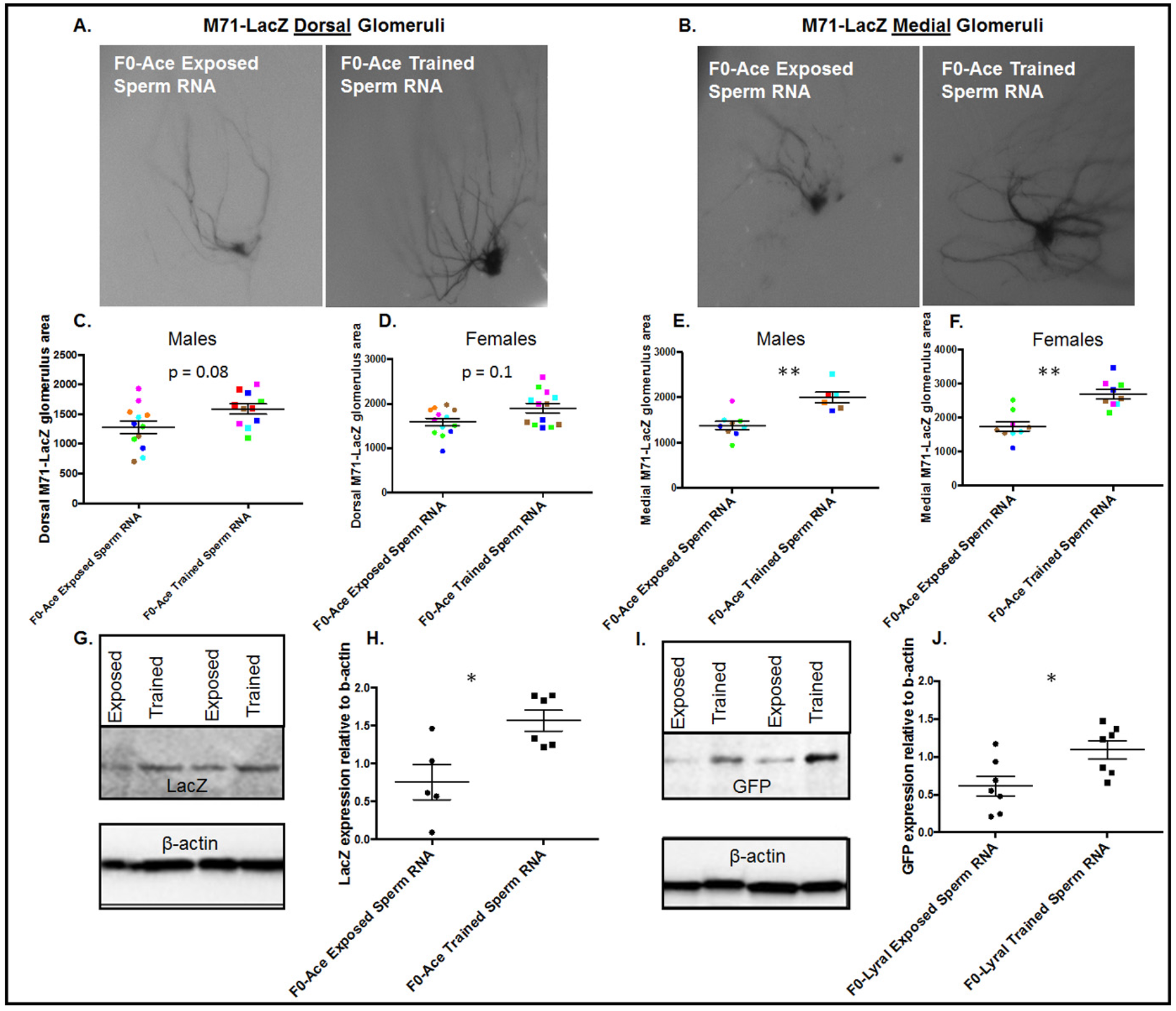
Sperm RNA contribute to neuroanatomical imprints of salient olfactory experiences across generations. **(A-F) β-galactosidase staining** shows that mice that developed from zygotes into which RNA from sperm of F0-Ace Trained males had been injected (F0-Ace Trained Sperm RNA) have larger dorsal and medial M71-LacZ Ace-responsive glomeruli compared to mice that developed from zygotes into which RNA from sperm of F0-Ace Exposed males had been injected (F0-Ace Exposed Sperm RNA) (F0-Ace Exposed Sperm RNA: Dorsal Glomeruli - Males: n = 12, 6 litters, Females: n = 13, 6 litters. Medial Glomeruli - Males: n = 11, 5 litters, Females: n = 9, 5 litters) (F0-Ace Trained Sperm RNA: Dorsal Glomeruli - Males: n = 11, 6 litters, Females: n = 13, 5 litters. Medial Glomeruli - Males: n = 6, 4 litters, Females: n = 9, 5 litters). Colors denote animals from a litter. **(G,H) Quantitation of LacZ expression in the MOE using Western Blotting.** F0-Ace Trained Sperm RNA animals (n = 6, 6 litters) have higher M71-LacZ expression in the MOE compared to F0-Ace Exposed Sperm RNA animals (n = 5, 5 litters). **(I,J) Quantitation of GFP expression in the MOE using Western Blotting.** F0-Lyral Trained Sperm RNA animals (n = 7, 7 litters) have higher MOR23-GFP expression in the MOE compared to F0-Lyral Exposed Sperm RNA animals (n = 7, 7 litters). Data presented as Mean ± SEM. * p < 0.05, ** p < 0.01.

#### F0-Lyral treatment

Due to the position of MOR23-GFP glomeruli on the olfactory bulb it is challenging to visualize and quantitate their size and instead we used western blotting to measure the GFP levels as has been reported [34] and as we have done in our previous publication [35]. We found a significant increase in GFP expression in the MOE of F1-Trained Lyral F0 Sperm RNA animals compared to F1-Exposed Lyral F0 Sperm RNA animals (t=2.681, df = 12, p< 0.05, Cohen’s D = 1.4) (Fig. 4I,J).

### Female offspring of F0-OdorA-Trained males show enhanced consolidation of weak singletrial Odor A+ shock fear conditioning

F0-Ace treatment: F0 males were either conditioned or merely exposed to Acetophenone. Their offspring were then fear conditioned to Acetophenone using a weak single-trial fear conditioning protocol wherein a single Acetophenone presentation was paired with a mild foot-shock in Context A. Their memory of this event was then measured in another context by measuring their freezing responses to an Acetophenone presentation in Context B. F1-Ace Trained female, but not male mice showed increased freezing to the Acetophenone presentation in Context B compared to F1-Ace Exposed animals (Female mice: t=2.13, df = 15, p<0.05, Cohen’s D = 0.87. Male mice: t=0.48, df = 8, p=0.643, Cohen’s D = 0.46) (Fig. 5A,B).

**Fig. 5:**
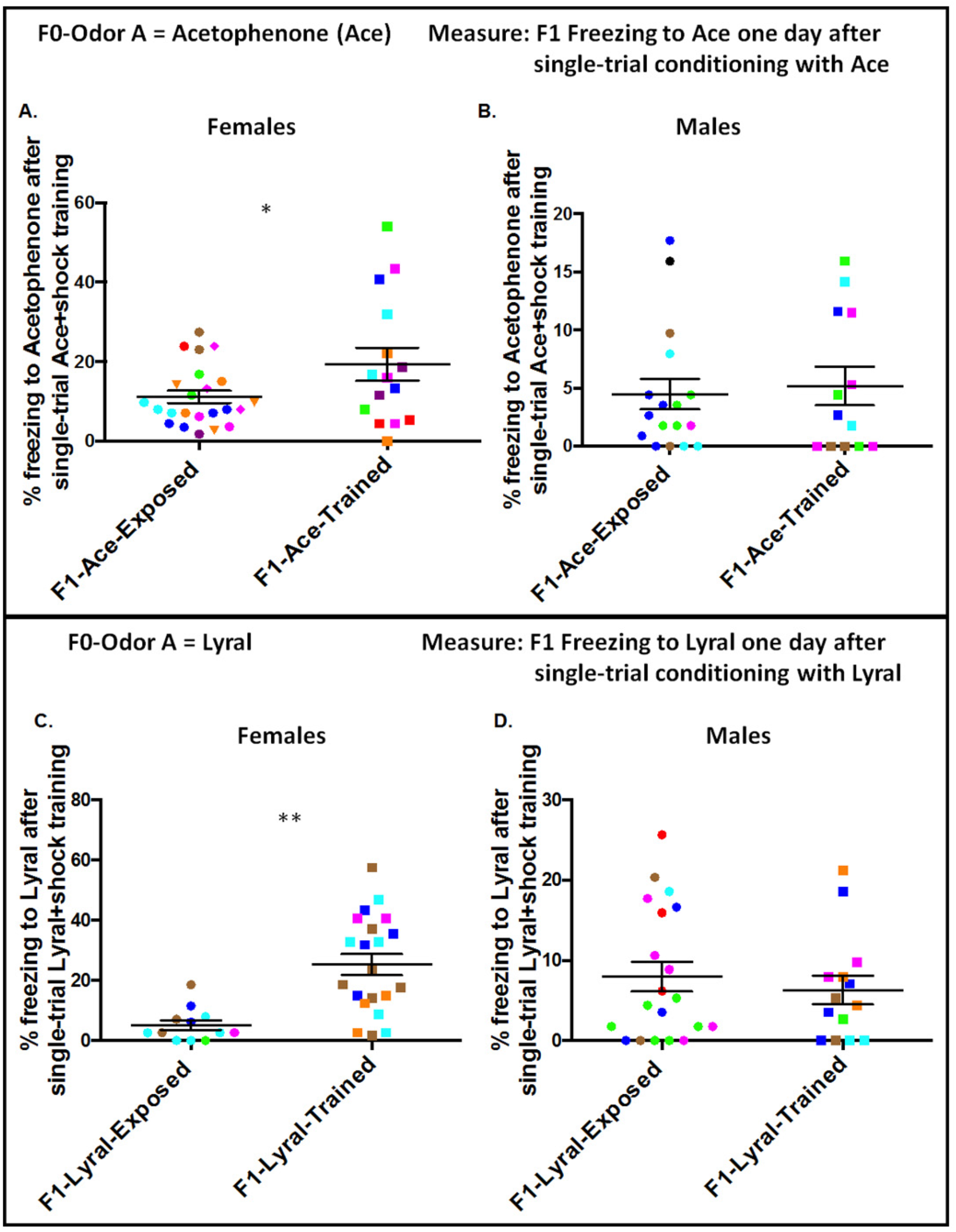
Parental sensory experience facilitates sensory learning in offspring. (A,B) Salient experience with Ace facilitates sensory learning about Ace in female offspring. Female **(A)** but not male **(B)** offspring sired by F0-Ace Trained males (F1-Ace-Trained) show higher consolidation of single trial Ace+shock conditioning compared to female offspring sired by F0-Ace Exposed males (F1-Ace-Exposed) (F1-Ace-Exposed: Males n = 17, 5 litters. Females n = 23, 10 litters) (F1-Ace-Trained: Males n = 13, 5 litters. Females n = 15, 7 litters). **(C,D) Salient experience with Lyral facilitates sensory learning about Lyral in female offspring.** Female **(C)** but not male **(D)** offspring sired by F0-Lyral Trained males (F1-Lyral-Trained) show higher consolidation of single trial Lyral+shock conditioning compared to female offspring sired by F0-Lyral Exposed males (F1-Lyral-Exposed). (F1-Lyral-Exposed: Males n = 20, 6 litters. Females n = 12, 5 litters) (F1-Lyral-Trained: Males n = 15, 5 litters. Females n = 21, 5 litters). Data presented as Mean ± SEM. Colors denote animals from a litter. * p < 0.05, ** p < 0.01.

F0-Lyral treatment: F0 males were either conditioned or merely exposed to Lyral. Their offspring were then fear conditioned to Lyral using a weak single-trial fear conditioning protocol wherein a single Lyral presentation was paired with a mild foot-shock in Context A. Their memory of this event was then measured in another context by measuring their freezing responses to a Lyral presentation in Context B. F1-Lyral Trained female, but not male mice showed increased freezing to the Lyral presentation in Context B compared to F1-Lyral Exposed animals (Female mice: t=4.19, df = 8, p< 0.01, Cohen’s D = 2.54. Male mice: t=0.43, df = 9, p=0.6751, Cohen’s D = 0.24) (Fig. 5C,D).

## DISCUSSION

Injecting RNA extracted from sperm of male mice that had been exposed to a salient olfactory experience like olfactory fear conditioning (Odor A+footshock) into single cell zygotes resulted in enhanced sensitivity for Odor A and increased representation of Odor A-responsive olfactory neuroanatomy in adulthood. F1 female, but not male mice exposed to weak single-trial olfactory conditioning with the odor that the F0 generation had been conditioned (Odor A) showed enhanced consolidation of single-trial Odor A+footshock conditioning. Using the framework provided by olfactory fear conditioning, our data provide evidence that intergenerational influences of salient sensory experiences can be mediated in part by RNA found in the parental male germline, and one consequence of such influence is to bias future learning about the sensory cues with which the parental generation had a salient experience.

Across species, experiences with sensory stimuli influence neuroanatomy, gene expression, physiology and behavior in the organism directly exposed to salient sensory stimuli [36–38]. It is becoming increasingly appreciated that the effects of parental exposure to salient sensory stimuli extend into offspring. Many cases of such intergenerational influences of sensory environments arise via social transmission through maternal behavior [12] and via the transfer of chemical cues across maternal-fetal barriers or egg barriers in viviparous or oviparous species, respectively [8, 9, 39]. While there is accumulating evidence that intergenerational influences of sensory environments are observed in offspring of worms, fruit flies and mice that were not conceived at the time of parental exposure to salient sensory stimuli [22–25, 37], mechanisms underlying these influences are unknown. To our knowledge, our study is the first to demonstrate that RNA found in the male germline of mammals exposed to salient olfactory experience can contribute to impacting olfactory-related neuroanatomy and function in future generations. Given that RNA contained in sperm have been demonstrated to be important contributors to intergenerational influences of paternal dietary manipulations and paternal stress exposure in rodents [26, 27, 29, 30], our work provides further evidence for germline RNA playing an important role in intergenerational influences of salient paternal environments, and novel evidence for their role in intergenerational influences of sensory experiences. A variety of RNA species in sperm like tRNA-fragments [26, 30], microRNA [29] and long RNA [28] have been demonstrated to orchestrate intergenerational influences of stressors and our future efforts will need to address which specific RNA in sperm are responsible for the intergenerational influences of salient sensory experiences.

Intergenerational influences of salient parental environments typically conjure up the idea that they place offspring at risk for neuropsychiatric disorders and an inability to cope with environmental challenges [40–42]. In contrast, a contrasting view is also developing in that intergenerational influences of salient parental environments can confer adaptive flexibility to the offspring that enables them to thrive in the face of environmental challenges [43]. Stress exposure to a parental generation of mice confers behavioral flexibility to offspring across a variety of behavioral tasks [44]. High levels of cocaine self-administration in rats confers upon the next generation, a lower propensity to abuse cocaine [45]. Raising gravid female crickets in a predator-rich environment results in the offspring showing an avoidance of predator-related cues in adulthood [39]. In keeping with this sentiment, our data also suggest an adaptive element to the behavioral sensitivity of F1 offspring to Odor A that we have previously demonstrated [22, 23]. More specifically, our data suggest that the pre-existing sensitivity to Odor A (that we have refrained from calling fear toward Odor A in this and our previous publications) confers upon the F1 (female) offspring the ability to learn about the salience of this odor even in a context that would not promote learning. Why we observe effects only in female offspring in this task and not in male offspring is an open question for which we do not have any definitive answers and can only offer speculation. It is not uncommon to observe sex-differences in the effects of intergenerational influences of salient parental environments. Exposing parental rodents to stressors as well as dietary manipulations have been shown to affect one sex and not the other depending not only on the parental perturbation but also the task on which the offspring are tested [46–53]. Therefore, once again, our data agree with the existing literature that male and female offspring may shoulder the legacy of parental stress differently and attention needs to be paid to the parental environmental experience and the dependent variable being tested in the offspring. With specific relevance to olfaction, sex differences in olfactory acuity have been reported [54–56] and our observation that female but not male F1 offspring show enhanced learning about singletrial conditioning with the odor used to condition the F0 generation may be a consequence of this sex difference. In the future, we will examine whether brain regions that respond to olfactory stimuli and that are important for associative learning like the piriform cortex and basolateral amygdala, respectively, are more active in F1 single-trial Odor A-conditioned female offspring exposed to Odor A than their F1 male conspecifics.

While we demonstrate that RNA contained in sperm of Odor A-trained F0 male mice renders adults generated from embryos into which they are injected, more sensitive to Odor A and possessing enhanced neuroanatomy for Odor A, we do not know if there is a general enhanced sensitivity to other odors (e.g. Odor B) or enhancements in olfactory neuroanatomy that respond to Odor B. Additionally, we do not yet know if F1 offspring sired by F0 males that have been conditioned with Odor A, also show a bias in their ability to consolidate memories of weak olfactory fear conditioning to Odor B. Such studies will be important future directions to determine whether the proximate causes and consequences of intergenerational influences of salient parental sensory experiences on offspring are specific to the parental sensory experience or whether there is a generalizability to a broader bouquet of sensory cues.

In summary, our data provide novel evidence that RNA in the male germline can contribute to intergenerational influences of parental sensory experiences and that one consequence of this phenomenon is allowing for adaptive responses of offspring to sensory cues when these cues are encountered under salient conditions. Our work fits into the broader landscape of studies that examine intergenerational influences of parental experiences and demonstrates that proximate mechanisms (RNA found in sperm) and consequences (adaptive responses) are conserved across parental experiences as diverse as stress exposure, dietary perturbations and sensory experiences.

## ACKNOWLEDGEMENTS

We thank the Veterinary and Animal Care staff in the Yerkes Neuroscience Vivarium for animal husbandry. Funding for this study was provided to BGD by the Emory University Department of Psychiatry and Behavioral Sciences, the Emory Brain Health Institute, the Yerkes National Primate Research Center (YNPRC), a CIFAR Azrieli Global Scholar Award and the Catherine Shopshire Hardman Fund. Additional funding was provided to YNPRC by Office of Research Infrastructure Programs ODP51OD11132.

## AUTHOR CONTRIBUTIONS

HSA, SS, SCH and ND performed experiments and analyzed data. HW analyzed data. AC provided technical feedback on intra-zygotic RNA injections. BGD designed the study, analyzed data, interpreted data and wrote the manuscript.

## AUTHOR INFORMATION

Correspondence and requests for materials should be addressed to.

